# A Ligand-Centered Framework for γδ T Cell Activation in Colorectal Cancer Revealed by Single-Cell and Transformer-Based Perturbation

**DOI:** 10.1101/2025.06.16.660020

**Authors:** Ran Ran, Douglas K. Brubaker

## Abstract

Understanding the activation mechanisms of γδ T cells in colorectal cancer (CRC) is critical for harnessing their therapeutic potential. Here, using an atlas of human CRC-infiltrating γδ T cells that we built by integrating multiple single-cell RNA-seq datasets, we developed a γδ T cell-refined ligand inference pipeline by combining differential gene expression, gene regulatory network prediction, ligand inference, and in silico perturbation analysis. This approach identified IL-15 and TNFSF9 (4-1BBL) as candidate ligands promoting γδ T cell effector function, and highlighted NCR2 and KLRC3 (NKG2E), whose in silico overexpression was associated with γδ T cell activation. Ligand enrichment analyses further indicated that monocytes and dendritic cells are key contributors to γδ T cell activation within the tumor microenvironment. Together, our results offer a systems-level view of the signaling and transcriptional programs governing γδ T cell phenotypes in CRC and provide a foundation for γδ T cell-based immunotherapies with enhanced antitumor function.

## INTRODUCTION

Colorectal cancer (CRC) is the one of the most frequently diagnosed cancer and the leading cause of cancer deaths^1–3^. It originates in the colon or rectum and is characterized by the abnormal proliferation of epithelial cells^4^. Surgery is the primary treatment for advanced resectable CRC, yet recurrence occurs in a substantial proportion of patients with stage II or III disease^5,6^. Most of recurrence occur within 5 years after surgery, and the relapse significantly increases the risk of metastasis and decreases the survival rate^5,7,8^. For non-resectable CRC, radiotherapy and chemotherapy are standard-of-care therapies that can reduce tumor burden^9,10^ but are limited by systemic toxicity, adverse effects, and lack of tumor specificity^11^. A substantial proportion of patients eventually develop incurable recurrent disease, even after standard treatments^11^. Therefore, there is a need for novel treatments that can be used in addition to conventional treatments to prevent recurrence.

γδ T cells are a unique subset of immune cells that have potent anti-tumoral ability^12,13^. They can be activated independently of classical MHC-mediated antigen presentation, and in some cases, through TCR-independent mechanisms involving engagement of stress-induced ligands on target cells^14^. Although γδ T cells only account for less than 5% of human blood T lymphocytes, their proportion is significantly higher in the colon mucosa^15^. They account for 10-40% of T cells in the epithelium monolayer^16–18^, where the CRC oncogenesis usually begins^19^. These cells exhibit dynamic surveillance behavior, migrating through the intestinal epithelium in a flossing-like manner and traversing the basement membrane to access the lamina propria^15,20^. γδ T cells have been shown to inhibit intestinal tumor growth via CD103-E-cadherin interactions, highlighting their role in maintaining epithelial integrity and tumor surveillance under homeostatic conditions^21^. Therefore, they are promising candidates for immunotherapeutic strategies because they are naturally present in the tumor microenvironment, exhibit strong tissue infiltration, and recognize targets in an MHC-independent manner, enabling potential allogeneic transfer.

However, despite its promise, the activation mechanism of γδ T cells in CRC remains unclear. In our previous work, we integrated single-cell RNA sequencing (scRNA-seq) data from multiple CRC studies and revealed diverse states among CRC-infiltrating γδ T cells. These included a substantial population of unexhausted, tissue-resident bystanders expressing *ITGAE* and *ITGA1*, as well as highly cytotoxic effector cells expressing *IFNG*^22^. We therefore hypothesize that identifying the molecular cues driving the activation of these bystanders could support tumor cell elimination and reduce CRC recurrence, given that recurrence often arises from residual tumor cells^7^.

It is well established that intraepithelial γδ T cells interact closely with colon epithelial cells under homeostatic conditions^21^. Through both T cell receptor (TCR)-dependent and TCR-independent pathways, they detect ligands expressed on epithelial cells or present in the local microenvironment. T cell activation also depends on the co-stimulatory ligands they engage and the cytokines they receive. Taken together, these insights support the hypothesis that γδ T cells may undergo a shift from a less activated to a more activated state through TCR-independent ligand-receptor interactions.

Single-cell RNA sequencing is transforming CRC research by enabling high-resolution profiling of gene expression in heterogeneous tumor tissues, capturing the activity of individual cell types including tumor-infiltrating immune cells. Here, we combine our previous multi-study integration of γδ T cell scRNA-seq data with gene regulatory network (GRN) inference and large language model (LLM)-based in silico gene perturbation to uncover the molecular drivers of γδ T cell activation in CRC. Through this integrative approach, we identify genes that correlate with—and may potentially drive—the activation transition, and make backward inferences about upstream ligands regulating them via signaling pathways.

## RESULT

### CRC-Infiltrating γδ T Subsets Have Varied Activation Levels

In our previous work, we integrated γδ T cells from multiple human CRC datasets to construct a comprehensive cell atlas that has over 18,000 γδ T cells—where 9,201 of them are identified as isolated from tumor by the data providers—across diverse anatomical sites, patient demographics, and disease stages, providing a broad view of γδ T cell transcriptional states in CRC^22^ (Fig. 1a). We performed clustering on this integrated dataset and identified γδ T cell subtypes based on canonical αβ T cell functional subset marker genes curated from the literature, including effector T cells (Teff), tissue-resident memory T cells (TRM), progenitor exhausted T cells (Tpex), and exhausted T cells (Tex), all present in both tumor and adjacent normal tissues, with Teff and Tex more enriched in tumors. We observed that the TRM-like population—the major subtype among tumor-infiltrating γδ T cells in our integrated dataset (Fig. 1b)—did not highly express conventional exhaustion markers such as *PDCD1* and *TIGIT*, nor effector-related genes including *GZMB*, *NKG7*, *PRF1*, or *IFNG* (Fig. 1c). Gene set scoring based on effector genes (*IL2RA*, *CD38*, *NKG7*, *GZMB*, *PRF1*, *IFNG*) and exhaustion genes (*PDCD1*, *CTLA4*, *LAG3*, *HAVCR2*, *TIGIT*) further confirmed that the TRM-like population exhibited the lowest scores for both effector and exhaustion signatures (Fig. 1d).

**Figure 1.**
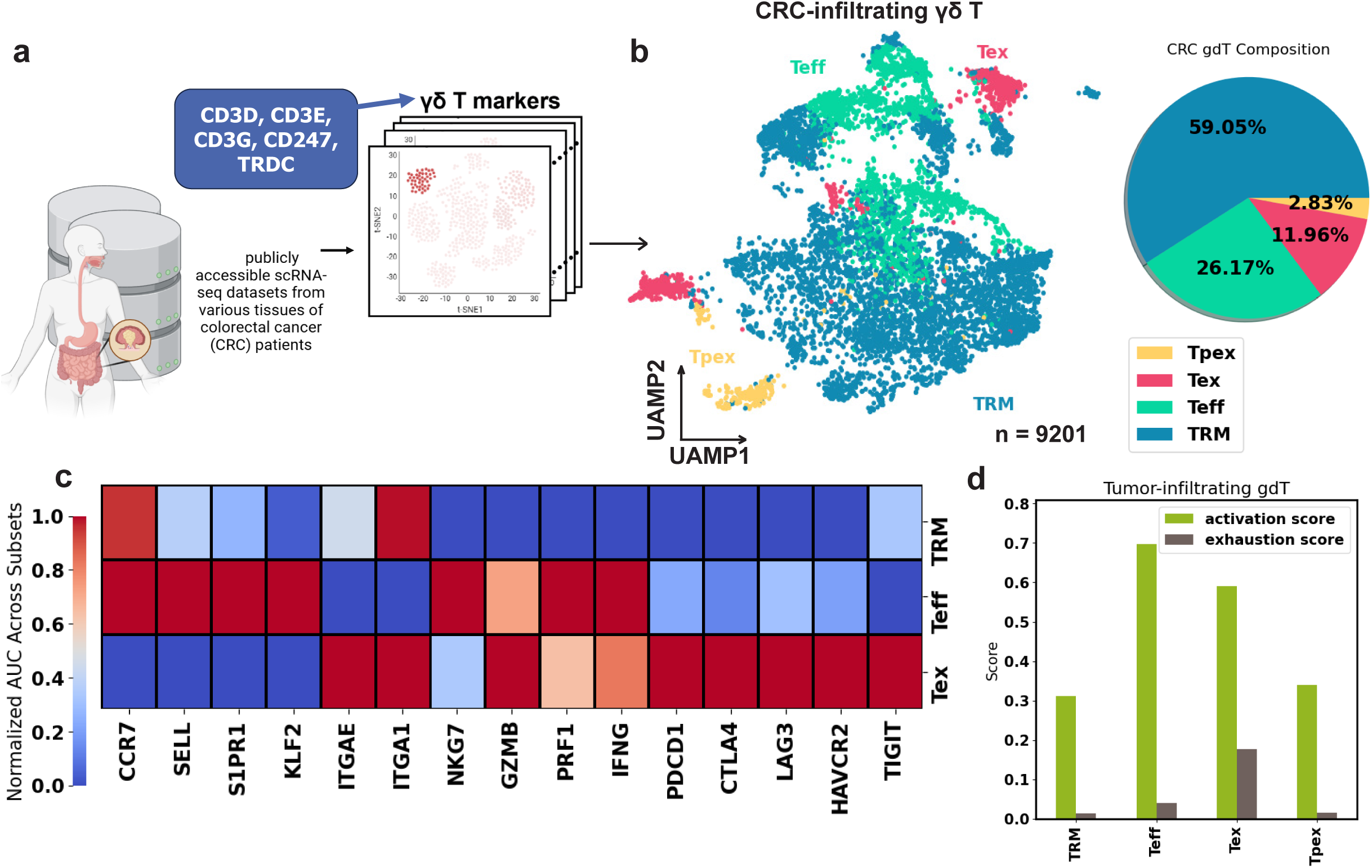
Overview of the heterogenous human CRC γδ T cells. (A) Using the minimum set of γδ T cell transcriptomics markers we developed, we examined publicly available CRC scRNA-seq datasets and identified γδ T clusters. They are integrated to build a human CRC γδ T cells dataset. (B) (Left) Uniform manifold approximation and projection (UMAP) of γδ T cells showing subsets: Teff (effector-like T cells), TRM (tissue-resident memory T cells), Tpex (progenitor exhausted T cells), and Tex (exhausted T cells). (Right) Composition of γδ T cell subtypes in the tumor. (C) Heatmap showing the T cell function related marker RNA expression in TRM, Teff, and Tex. Expression has been normalized across cell types. (D) Bar plot showing the effector activation score and exhaustion score in γδ T cell subsets.

Given the plasticity of T cells, we aimed to identify candidate ligands that could drive the potential transition from TRM-like bystanders to Teff cells in CRC. Using scRNA-seq data, we developed a pipeline that integrates multiple computational tools to perform ligand inference (Fig. 2). At the core of this pipeline is NicheNet^23^, a computational framework that performs random walk on a ligand to target gene transcription network that is constructed from known ligand receptor interactions, intracellular signaling, and gene regulatory network to infer the ligands that are most influential in regulating a given set of genes. For the standard processing, we input differentially upregulated genes in the Teff compared to TRM as the signature associated with the state transition into NicheNet, which will output the ligands that most influence these differential expression events.

**Figure 2.**
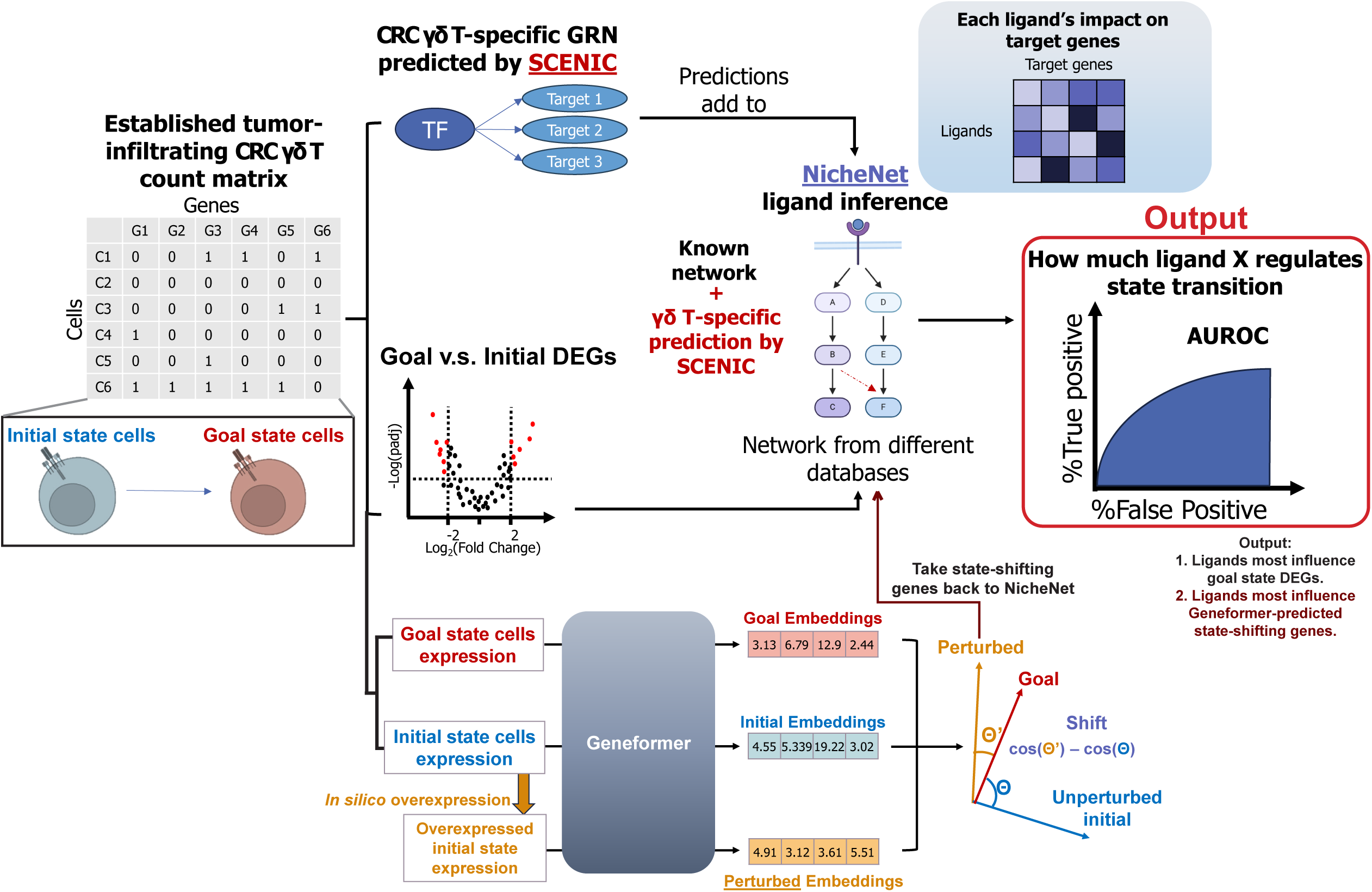
γδ T cell-refined ligand inference pipeline. Overview of the computational pipeline. We first define a potential cell states transition from an initial state (i.e., Teff) to a goal state (i.e., TRM) in the scRNA-seq count matrix. Then, we use NicheNet, a ligand inference tool that uses Personalized Page Rank on the constructed ligand to target gene network—known networks summarized by NicheNet authors from databases and γδ T- specific networks predicted by SCENIC based on the co-expression of known TF with potential target genes in our data—to approximate the propagation of the signal in order to compute each ligand’s regulatory power on its target gene transcription. By computing the AUROC between the transition-associated signatures and each ligand to target genes regulatory potential vector, we can infer which ligand has the highest potential of regulating the given signatures. The transition signatures are calculated in two ways: differential gene expression analysis, and in silico perturbation analysis using Geneformer, a language model.

One challenge in this style of ligand inference is that the differentially expressed genes (DEGs) between TRM and Teff γδ T cells identifies genes associated with their transcriptional differences, but such differences reflect a mixture of upstream, downstream, and unrelated processes. To prioritize gene programs more likely to contribute to the transition between these states, we incorporated Geneformer^24^, a large language model pre-trained on human scRNA-seq data for in-silico perturbation analysis, into our pipeline to identify genes whose in silico overexpression is associated with a shift from a TRM-like to a Teff-like transcriptional profile. In our modified workflow, NicheNet is used to calculate another set of ligands that can regulate the Geneformer-predicted genes expression events leading to the cell state transition.

A further challenge in ligand signaling inference is that gene regulatory networks (GRN) are cell type-specific^25^ and γδ T cells are a less well-studied population whose cellular programs may not be well-represented in prior knowledge networks. To address this limitation of existing workflows, we will integrate ligand-receptor interactions relevant to γδ T cells, curated from the literature,^14,26–31^ with SCENIC^32^-inferred transcription factor (TF)-target gene relationships, into the NicheNet signaling network and GRN by adding the γδ T cell-specific edges to NicheNet’s current default network. Using this γδ T cell-tailored network, we can perform finely tuned γδ T-specific ligand inference to prioritize candidate ligands and gene expression programs associated with γδ T cell activation in CRC.

### Integration of SCENIC-Predicted Novel Gene Regulatory Relationships Maintains General Predictive Performance

The first step was to infer CRC-infiltrating γδ T cell-specific TF-target gene relationships from our integrated single-cell atlas. Using SCENIC, we identified 30,025 significant and distinct TF-target gene pairs in our γδ T cell atlas, of which 56% were already documented in NicheNet’s default GRN (Fig. 3a; full list of regulons in Table S1). We used publicly available human T and NK cell ChIP-seq data to assess whether the remaining 44%—approximately 13,200 novel TF-target gene pairs specific to CRC-infiltrating γδ T cells—could be validated in an orthogonal experimental setup. From ChIP-Atlas^33–35^, 19 of the predicted TFs had available peak information, and for 15 of them, ChIP-seq peaks were found within ±5 kb of the transcription start site (TSS) of at least a subset of their SCENIC-predicted target genes (Fig. 3b). Regulon specificity scores across the four γδ T cell subsets indicated distinct TF regulatory programs, with TRM cells primarily regulated by TFs such as IKZF1 and FOXO1, Teff cells by KLF2 and interferon regulatory factor 1 (IRF1), Tex cells by MYBL2 and SPI1, and Tpex cells by RUNX1, EGR3, and NFKB1 (Fig. 3c).

**Figure 3.**
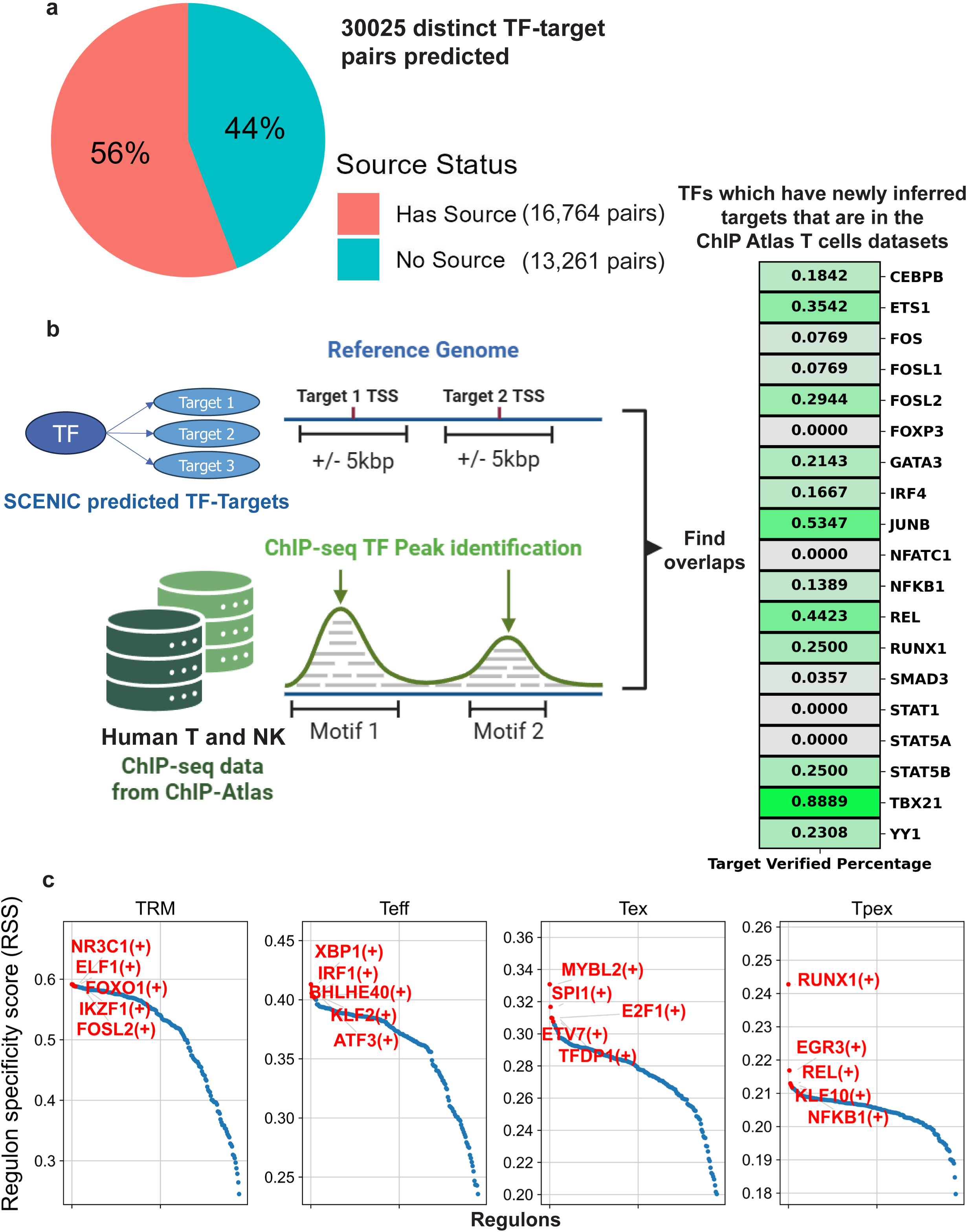
SCENIC Predicts Gene Regulatory Networks and TF Activities in Human CRC γδ T cells. (A) Pie chart showing the percentage of SCENIC predicted TF-target gene pairs are already in NicheNet-summarized GRN. (B) (Left) Workflow of using ChIP-seq data to validate SCENIC prediction. Overlap between a TF’s target genes TSS surroundings and that TF’s ChIP-seq peaks, if contained in the database, is calculated. (Right) Heatmap visualizing the percentage of TF’s SCENIC-predicted target genes can be found to overlap with a peak. (C) The regulon specificity score plot showing how the collective expression of a TF’s target genes in cells converge with the cell type labels, or in other words, how specific a TF’s regulatory activities is to the cell type. The x-axis is the ranked regulons.

All novel TF-target gene pairs were incorporated into NicheNet’s gene regulatory network. In the original NicheNet study, model performance was evaluated by predicting ligands from RNA-seq data derived from experimentally perturbed conditions using known ligands, referred to as the “gold standard.” Because the γδ T cell-specific additions primarily expand underrepresented signaling and regulatory interactions rather than altering shared core pathways, we expected model performance on unrelated test data to remain stable. Indeed, benchmarking confirmed that integrating both the SCENIC-predicted GRN and literature-curated γδ T cell ligand-receptor interactions did not compromise the model’s ability to correctly identify perturbed ligands in the reference dataset (Fig. S1).

### NicheNet Combined with SCENIC Infers Ligands Including 4-1BBL and IL-15 That Shape γδ T Cell Activation in CRC

After customizing the NicheNet model, we computed DEGs between Teff and TRM γδ T cells in our atlas. In addition to classical cytotoxic-molecule encoding genes such as *GZMB*, *NKG7*, and *GNLY*, and effector function-related genes like *IFNG*, Teff cells also upregulated *KLRG1*, *CX3CR1*, and *ZNF683* (encoding Hobit) (Fig. 4a; full list in Table S2). Collectively, these DEGs reflect the effector-associated transcriptional profile of Teff cells.

**Figure 4.**
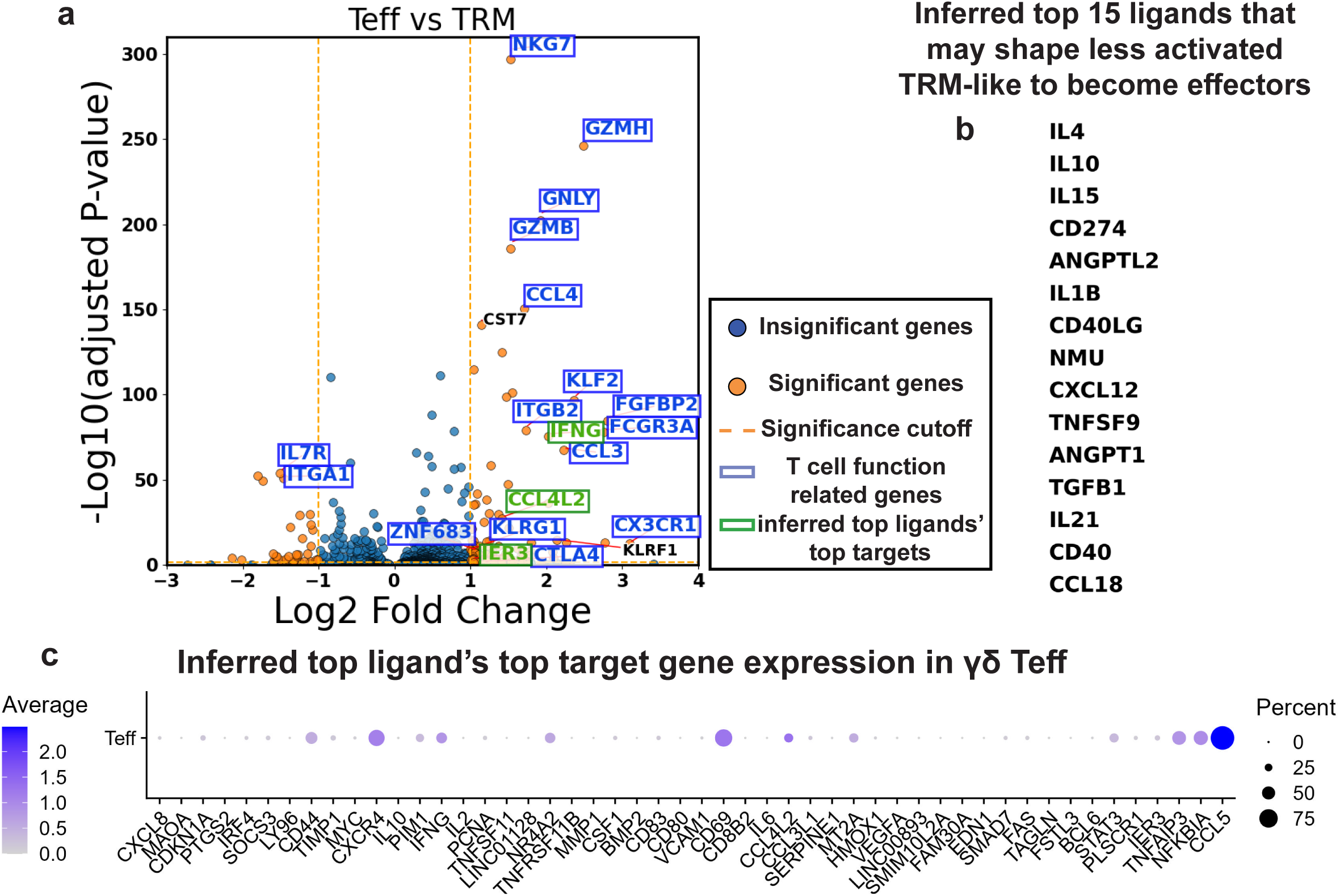
NicheNet Predicts Ligands that Most Influence the TRM to Teff Transition from Differential Gene Expression. (A) Volcano plots showing the differentially expressed genes between Teff and TRM (Teff upregulation on right side). Each dot is a gene. X-axis shows the log2 fold change in gene expression between two groups, and the y-axis shows the -log10 of multi-comparison-corrected P-value of each gene’s differential expression being statistically significant. (B) Top (ranked by area under the receiver operating characteristic curve (AUROC)) NicheNet-inferred ligands that shape the differential upregulation in Teff with respect to the TRM, colored by the AUROC value. (C) Dot plot showing the expression and fraction of expressing cells of the top 15 NicheNet-inferred ligands’ top 5 target genes in Teff cells.

To identify potential upstream regulators of this transcriptional program, we input the Teff-upregulated genes into NicheNet to infer ligands most likely to drive the TRM-to-Teff transition. Among the top 15 predicted ligands were IL-4, IL-15, IL-21, and TNFSF9 (encoding 4-1BBL), suggesting they may contribute in promoting γδ T cell activation. However, the activation landscape also included immunosuppressive signals, with TGFB1, IL10, and PD-L1 (CD274) predicted as counter-regulatory ligands (Fig. 4b, list in Table S3). For comparison, we also used the DEG-based NicheNet approach to predict ligands potentially driving the transition from Teff to Tex. This analysis identified ligands corresponding to several inhibitory receptors, including CD274 (PD-L1, binding PD-1), LGALS9 (binding TIM-3), NECTIN2 (binding TIGIT), and CDH1 (E-cadherin, binding KLRG1), highlighting their potential roles in promoting effector exhaustion (Fig. 5a,b; list of DEGs in Table S4, list of ligands in Table S5).

**Figure 5.**
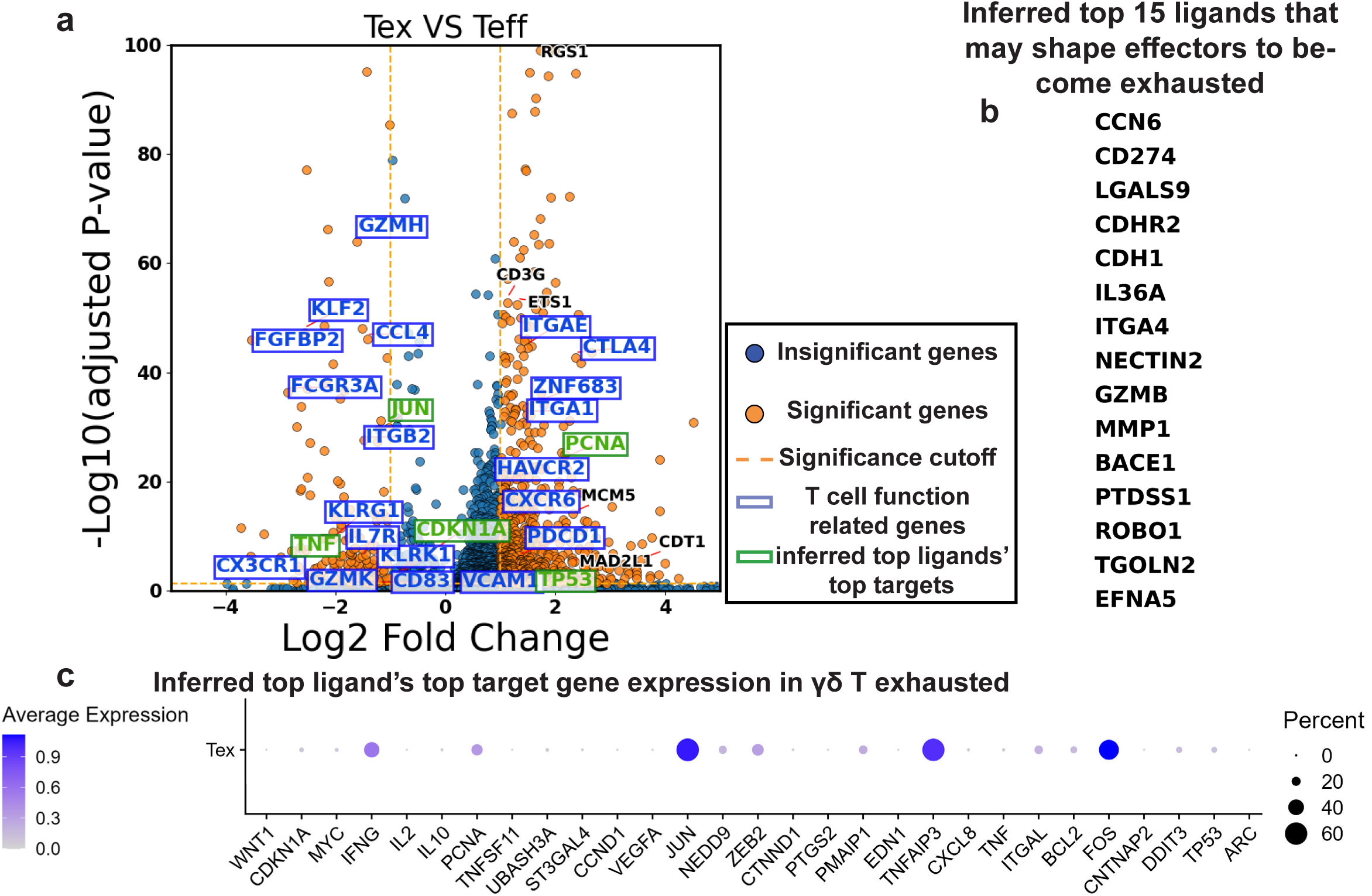
NicheNet Predicts Ligands that Most Influence the Teff to Tex Transition from Differential Gene Expression. (A) Volcano plots showing the differentially expressed genes between Tex and Teff (Tex upregulation on right side). Each dot is a gene. X-axis shows the log2 fold change in gene expression between two groups, and the y-axis shows the -log10 of multi-comparison-corrected P-value of each gene’s differential expression being statistically significant. (B) Top NicheNet-inferred ligands that shape the differential upregulation in Tex with respect to the Teff, colored by the AUROC value. (C) Dot plot showing the expression and fraction of expressing cells of the top 15 NicheNet-inferred ligands’ top 5 target genes in Tex cells.

Although NicheNet evaluates the full list of differentially expressed genes (DEGs) to generate ligand predictions, we asked which target genes were most strongly associated with the top-ranked ligands. Specifically, we examined the top five target genes for each of the top predicted ligands and assessed their expression in the goal cell states—Teff and Tex—to gain insight into the basis of the NicheNet predictions (ligand-target gene matrices in Fig. S2a,b). Several of the top target genes were indeed highly expressed in the goal state cells (Fig. 4c, Fig. 5c). Among them, *IFNG*, *CCL4L2*, and *IER3* stood out as some of the most upregulated genes in the Teff population (Fig. 4a).

We also compared the ligand prediction results from the unmodified NicheNet model with those from the γδ T cell-tailored NicheNet. The inclusion of γδ T cell-specific edges did not significantly increase the number of novel ligands predicted to perform better than random (defined as AUROC > 0.5), but it did adjust the relative ranking and confidence of ligand predictions within the γδ T cell context (Fig. S3a-c; full list of ligands predicted by unmodified NicheNet in Table S6, S7). This refinement can support future experimental design by prioritizing ligands that are more likely to have functional impact on γδ T cells. Using the annotated CRC whole-tissue scRNA-seq reference dataset from Joanito et al.^36^, we further assessed the enrichment of the top 30 γδ T cell phenotype-shifting ligands across various cell types within the CRC tumor microenvironment. iCMS2-type CRC cells, found in microsatellite-stable (MSS) patients, were enriched for ligands predicted to regulate the TRM-to-Teff transition, whereas iCMS3 cells, also from MSS patients, were enriched for ligands associated with a shift toward exhaustion (Fig. 6a; detailed expression heatmaps in Fig. S4a,b). In both MSS and microsatellite instability-high (MSI-H) CRC, monocytes/classical dendritic cells (McDCs) exhibited the strongest potential to drive γδ T cell transition from a less activated, TRM-like state to a more activated, effector phenotype (Fig. 6b; expression heatmap in Fig. S5a). Additionally, enrichment analysis indicated that CRC fibroblasts may contribute to the exhaustion of tumor-infiltrating γδ T cells (Fig. 6b; detailed ligand expression in Fig. S5b).

**Figure 6.**
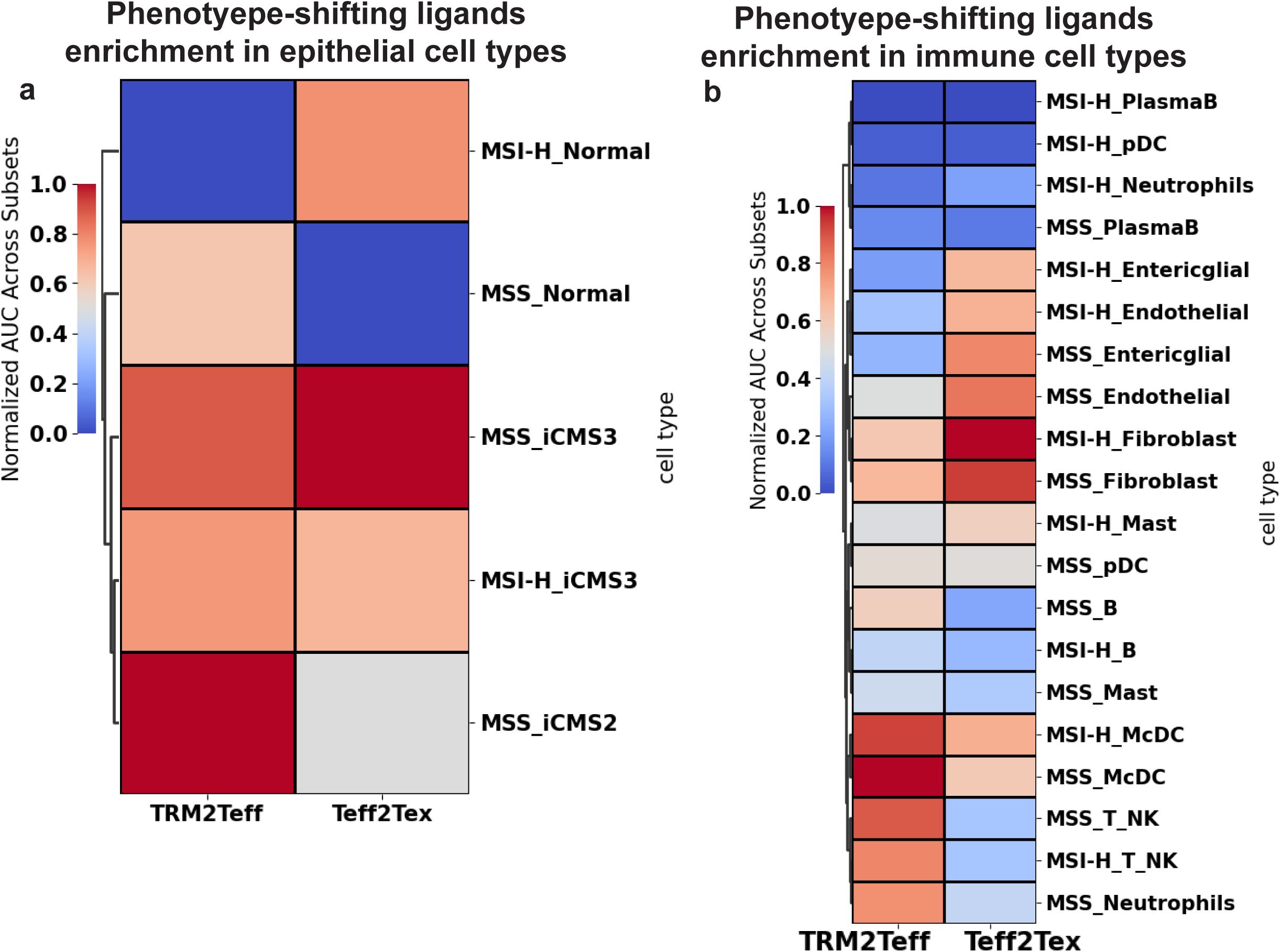
Human CRC γδ T cells Phenotype-shifting Ligands Are Enriched in Macrophage, Dendritic Cells, Fibroblasts, and iCMS2/3 CRC Cells. The enrichment of the NicheNet-inferred phenotype-shifting ligands in each CRC TME (A) epithelial (B) non-epithelial cell type were quantified by the area under the recovery curve (AUC) against the top 30 (ranked by area under the receiver operating characteristic curve (AUROC), AUROC > 0.5) NicheNet-inferred ligands. Results are visualized by heatmap, where the color represents the enrichment min-max normalized across cell types. TRM2Teff, TRM to Teff. Teff2Tex, Teff to Tex.

### Geneformer Predicts That Overexpression of *NCR2* and *NKG2E* Shifts Cell State Toward an Effector Phenotype

We next performed *in silico* perturbation on the TRM-like γδ T cell profile using Geneformer to identify which gene overexpression events would most effectively shift the cell state toward Teff. Briefly, Geneformer leverages a BERT-based masked language modeling (MLM) framework trained on over 95 million human single-cell expression profiles to learn gene-gene relationships across diverse biological contexts. This enables it to compute contextual embeddings of cell states based on their expression profiles. The *in silico* overexpression analysis involves simulating the increased expression of candidate genes within the original TRM cell state and assessing which gene perturbations produce a cell embedding that is similar to the embedding of the goal state—in this case, Teff (Fig. 2).

Using the fine-tuned Geneformer model—which achieved 88% accuracy in classifying TRM and Teff labels (Fig. S6)—we identified several genes whose in silico overexpression was associated with a shift in embedding toward an effector-like state. These included *CCR9*, *NCR2*, *KLRK1* (NKG2D), *KLRC3* (NKG2E), *NR4A3*, and *MMP9* (Fig. 7a, full list of DEGs in Table S8). Notably, most of these genes were not differentially expressed between Teff and TRM-like cells, indicating that Geneformer uncovered novel perturbation targets rather than simply retrieving genes whose differential expression reflects the Teff-like transcriptional profile. We then re-applied the γδ T cell-refined NicheNet to the set of Geneformer-predicted perturbation genes to infer upstream ligands. Among the top predicted ligands were IL-15 and TNFSF9, which also ranked highly in the initial NicheNet analysis based on differentially expressed gene signatures (Fig. 7b, full list of ligands in Table S9).

**Figure 7.**
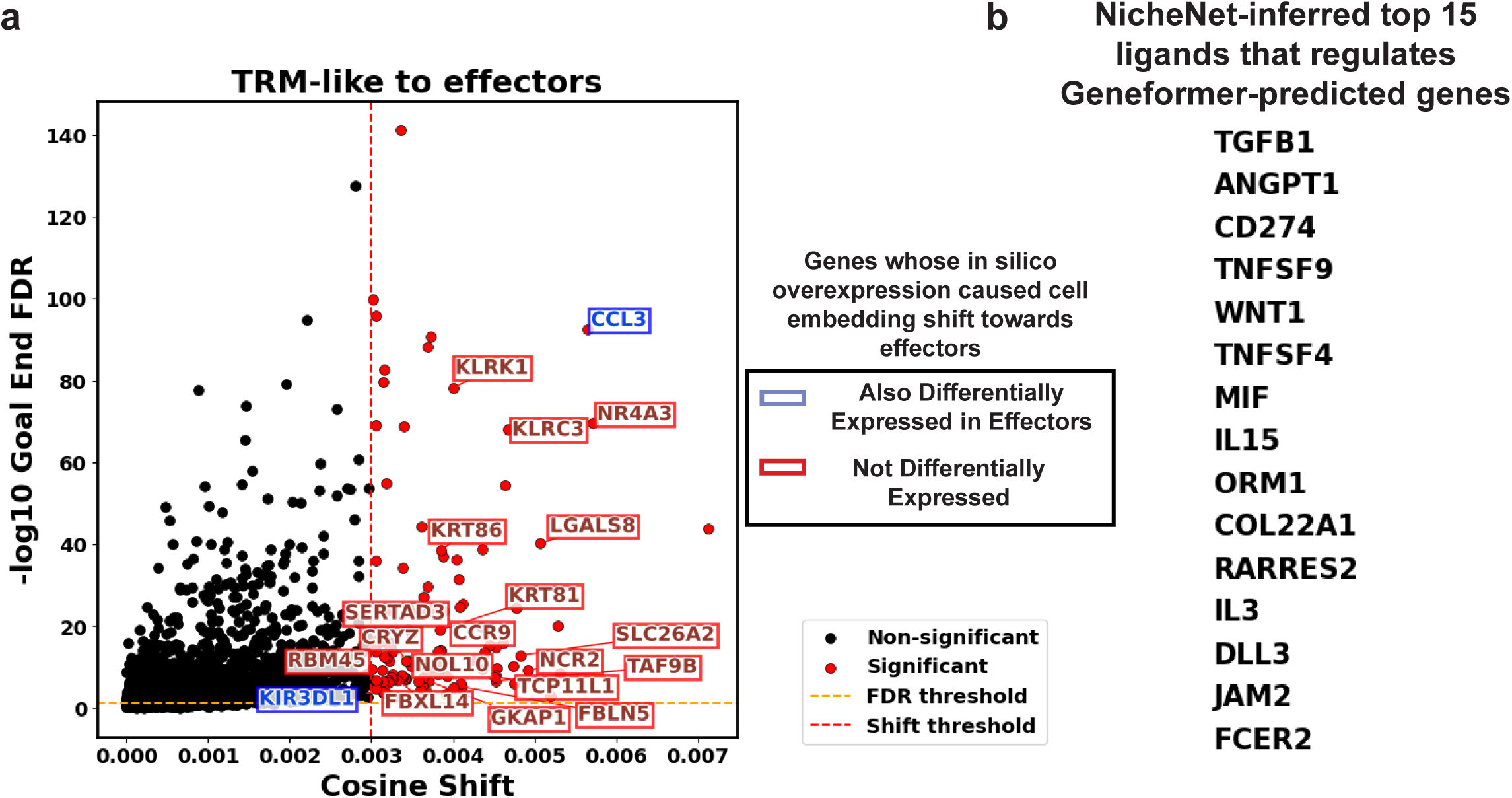
In Silico Perturbation Using Geneformer. (A) Scatter plot visualizing the cosine shift caused by the *in silico* overexpression of each gene from the TRM state towards the Teff state. X-axis shows the cosine shift, and the y-axis shows the multi-comparison-corrected P-value of each gene being statistically significant. (B) Top NicheNet-inferred ligands (derived from Geneformer-predicted state) that shape the differential upregulation in Teff with respect to the TRM, colored by the AUROC value.

## DISCUSSION

In this study, we developed a γδ T cell-tailored ligand inference framework to investigate the molecular drivers of γδ T cell activation in CRC. In our previous work, by integrating multi-study single-cell RNA-seq data, we constructed a comprehensive atlas of CRC-infiltrating γδ T cells and identified major subsets including TRM-like bystanders and cytotoxic effectors. To uncover signals responsible for driving transitions between these phenotypes, we combined differential gene expression analysis, SCENIC-inferred gene regulatory networks, NicheNet ligand inference, and the large language model Geneformer for in silico perturbation. We refined the NicheNet framework by incorporating γδ T cell-specific ligand-receptor interactions and transcriptional regulatory edges. This approach revealed key ligands—including IL-15 and TNFSF9—as potential regulators of γδ T cell activation. Notably, Geneformer identified perturbation targets such as NCR2 and KLRC3 (NKG2E), which were not differentially expressed but capable of inducing a Teff-like state, highlighting the importance of natural killer receptor signaling other than the well-known NKG2D and NCR3. Together, our framework provides a systems-level perspective on the extrinsic and intrinsic factors shaping γδ T cell phenotypes in CRC and offers a prioritized list of experimentally testable candidates for immunomodulation.

SCENIC revealed distinct TFs that regulate γδ T cell subtypes in human CRC. IKZF1(Ikaros), previously shown to repress gene programs associated with type I interferon responses^37^, may help explain the limited effector function observed in TRM-like γδ T cells. KLF2, whose downregulation is required for the establishment of tissue residency^38^, promotes *S1PR1* expression, facilitating T cell circulation through the bloodstream and lymphatic system^39–41^. It has also been implicated in enhancing effector function and preventing exhaustion in CD8⁺T cells during LCMV infection^42^, raising the possibility that KLF2 may influence effector function or exhaustion dynamics in γδ T cells. MYBL2, associated with cell proliferation^43^, may indicate a proliferative potential in γδ T cells expressing exhaustion markers. SPI1 acts downstream of PRDM1 to enhance PD-L1 transcription^44^, pointing to a regulatory role in the Tex γδ T cell subset.

Our DEG analysis highlights key features of γδ T effector cells in human CRC. *KLRG1*, a marker of terminal differentiation and senescence in CD8⁺ effector T cells^45,46^, was highly expressed in γδ Teff cells in our integrated dataset. As an inhibitory receptor, KLRG1 impairs T cell effector function by blocking AKT phosphorylation through the recruitment of phosphatases that convert PIP3 to PIP2^47^. In murine models, KLRG1 inhibition enhances antitumor activity^48^. Flow cytometry studies in human colorectal cancer have shown that KLRG1*⁺* T cells exhibit impaired cytotoxicity^49^; however, the functional consequences of KLRG1 expression in γδ T cells remain unclear. It is yet to be determined whether KLRG1 dampens γδ T cell function as it does in natural killer (NK) cells^50^, or whether its expression is more permissive—as has been debated for PD-1, which may not signify exhaustion in γδ T cells but instead reflect an activation state^51^.

Interestingly, KLRG1⁺ CD8⁺ Teff cells have been shown to downregulate KLRG1 and differentiate into memory subsets^52^. KLRG1⁺ γδ T cells are also found in CMV-uninfected newborns^53^ and become more prominent following CMV infection and in adulthood^54^. In murine cytomegalovirus (MCMV) infection, KLRG1⁺CX3CR1⁺ γδ effector memory cells populate lung and liver vasculature and mount strong antiviral responses^55^. Although a scRNA-seq analysis of CRC patient samples from TCGA reported enrichment of KLRG1⁺ cytotoxic T cell signatures associated with favorable outcomes^56^, it may be premature to attribute a positive role to KLRG1 in antitumor immunity.

Additionally, ZNF683 (Hobit), which has been implicated in promoting IFNG transcription in effector T cells^57^ and marks precursors of tissue-resident memory T cells^58^, was also upregulated in γδ Teff cells, supporting their cytotoxic potential and tissue-localized phenotype. Compared to the less activated TRM-like population, Teff cells also showed downregulation of *IL7R* and *ITGA1*, suggesting reduced emphasis on survival and increased migratory capacity.

Ligand prediction by NicheNet in the TRM-Teff transition provided many interesting targets for γδ TCR-independent activation. IL15, needless to say, is a well-studied ligand essential for T cell survival, proliferation, and effector function^59^. It has been used in multiple studies to expand Vδ2⁺ γδ T cells or CAR γδ T cells and enhance their cytotoxicity^60–62^, although there are also reports that continuous IL-15 signaling leads to exhaustion in NK cells^63,64^. Another interleukin in the list, IL21, is also known for promoting T cell proliferation and cytotoxic differentiation^65^. Literature has reported its effect in providing short-term proliferation^66^ and enhancing the antitumor cytotoxic response in γδ T cells in the presence of IL-2^67^. Studies also have shown IL-21’s synergistic effect with IL-15 in promoting CAR T and NK cell expansion^68,69^. Interestingly, both CD40 and its ligands, CD40LG, are in the top ligands list. Recent findings show that CD40LG expressed on CD8⁺ T cells can engage CD40 on dendritic cells, leading to their activation and promoting CD8⁺ T cell effector differentiation^70^. The CD40LG on CD8 T cells is found to interact with CD40 on dendritic cells, such that this interaction not only promote CD8 T cell’s effector function^71^ but also the survival of dendritic cells in antitumor response^72^. In mice parasite infection, γδ T cells enhance dendritic cells activation through CD40LG-CD40 interactions^73^. Considering that TRM-to-Teff NicheNet-predicted ligands were most enriched in McDCs in Joanito *et al.*’s dataset, a model where γδ T cells promote dendritic cell activation via CD40LG*-*CD40 interactions, thereby reinforcing their own effector programming, is a reasonable hypothesis. Indeed, although γδ cells are known for MHC-independent activation, our result of McDCs being identified as the top candidate influencing γδ cells effector differentiation encourages future studies that focus on antigen-presenting cells (APC)-γδ cells interactions.

On the other hand, there is also a possibility that CD40 is expressed on γδ T cells and interacts with CD40LG⁺ cells in the TME. Expression of CD40 on activated CD8⁺ T cells has been reported^74,75^, and in vitro stimulation of CD40⁺ human CD8⁺ T cells with CD40LG⁺ artificial APCs increased both the total number and percentage of effector cells^75^. Although such CD40 expression appears to be transient, it may still exert long-term effects on T cell differentiation, at least in terms of cell numbers as pointed out in the previous study^75^.

Another interesting finding is the presence of TNFSF9, the 4-1BB ligand, among the top Teff-shifting ligands, rather than CD80/86, which bind CD28. In T cells, TNFSF9 provides a potent co-stimulatory signal through 4-1BB, amplifying the proliferation and survival of effector T cells^76^. Since the second generation of CAR-T therapy, the intracellular domain of 4-1BB has been incorporated into the CAR design^77^ to achieve a better activation. Current CAR γδ T cells typically use Vδ2 TCRs, the most abundant γδ T subset in blood, and incorporate the CD28^78^ signaling domain based on second-generation CAR αβ T cell design^14^. However, whether CD28 effectively helps the activation of most intraepithelial γδ T cells, or generally speaking, the Vδ1 subset which has stronger cytotoxicity^79^, tissue residency^80^, and tumor infiltration^81–83^, remains unclear. Our preliminary scRNA expression data of intraepithelial γδ T cells in healthy human colon^84^ agrees with previous mouse^85^ and human^86^ observations that they don’t express CD28. Additionally, there are contradictory results of CD28 antibody’s effect on human γδ T cells^87,88^. There are multiple potential therapeutic co-stimulatory receptors like 4-1BB^89^, CD27^90^, and CD6^91^, but stimulating them in the CRC settings has unknown results, plus there could be more undiscovered. The identification of 4-1BBL among the top ligands but not CD80/86 suggests that 4-1BB is likely to serve as a more prominent or contextually relevant co-stimulatory signal for γδ T cells in human CRC. Indeed, ligand prediction based on Geneformer-derived overexpression targets also supported the importance of 4-1BBL.

Together, these findings highlight ligands such as IL-15 and TNFSF9 as promising candidates for promoting γδ T cell activation in CRC. In addition, NCR2 and NKG2E emerged as potential receptors through which γδ T cells may recognize CRC cells. These insights can inform the design of engineered γδ T cell therapies aimed at enhancing antitumor function and supporting both early tumor elimination and long-term post-surgical surveillance.

## METHOD

### Data Acquisition

The human CRC-infiltrating γδ T cell scRNA-seq dataset was curated as described in our previous work^22^. Briefly, we searched for publicly available scRNA-seq datasets from treatment naïve CRCs in indexed journals and Gene Expression Omnibus. In each study, cell clusters that highly co-expressed CD3D, CD3E, CD3G, CD247, and TRDC were identified as γδ T cells. Distinct γδ T cell clusters with more than 50 cells were identified from 9 studies that provided raw counts. Human CRC whole-tissue scRNA-seq datasets were obtained from Gene Expression Omnibus (GEO) Series GSE178341^92^, GSE200997^93^, GSE245552^94^, GSE231559^95^, GSE188711^96^, GSE161277^97^, GSE108989^98^, GSE178318^99^, and PubMed PMID35773407^36^.

### Data Processing

For processing the original dataset, only cells in the colon mucosa were kept. If processed data is not provided, Seurat^100^ and Scanpy^101^ were used for downstream analysis. For quality control, low-quality cells were dropped based on their low UMI counts (<500), high mitochondrial gene counts (>20%), and a low number of uniquely expressed genes (<200). If the unprocessed data of a study has multiple samples, data integration was done by using Seurat on the normalized and log-transformed raw counts, which means each cell’s counts are divided by its total counts, multiplied by 10,000, then log-transformed. SCTransform was applied to raw counts of γδ T cells isolated from each study. As we have briefly summarized in the previous work, raw counts were fitted into negative binomial distributions whose expectation was a function of the total count of cells. The coefficients of gene’s general linear model are further regularized by the kernel regression with the coefficient of other genes that have similar average expression. Regressed gene expression calculated from the regularized generalized linear model coefficients was referred as “SCTransform-corrected counts,” and log-transformed SCTransform-corrected counts were used for later visualization. The Pearson residual of a cell’s observed gene expression to the SCTransform-corrected counts was used for integration and later performing principal component analysis (PCA, 50 pcs kept). Leiden clustering was performed based on the computed neighborhood graph of observations (UMAP, 50 pcs, size of neighborhood equals 15 cells) to reveal the general subtypes. In order to delineate cell subtypes, further subclustering was performed on each subcluster at a resolution range from 0 to 1 (detailed resolution for each step of subclustering was recorded in the published code). The cell type annotating strategy was detailed in our previous work. In short, we identified four distinct subtypes of γδ T cells in our integrated dataset: poised effector-like T cells (Teff, markers: *KLF2, KLRG1*, *TBX21*, *IFNG* and low *IL7R, ITGAE, ITGA1, CCR7, SELL*), tissue-resident memory T cells (TRM, markers: *ITGAE, ITGA1* and low *CCR7, SELL, KLF2*), progenitor exhausted T cells (Tpex, markers: *TCF7, PDCD1, CTLA4, LAG3, TIGIT, HAVCR2*), and Tex (*GZMB*, *PDCD1, CTLA4, LAG3, TIGIT, HAVCR2* and no *TCF7*). Effector and exhaustion score were calculated for each cell based on the aforementioned genes using scanpy function score_genes against the same number (6) of random genes.

### SCENIC

Cell expression profile of the tumor-infiltrating γδ T cells was input into SCENIC (v1.3.1) pipeline using default parameters. List of TFs “hs_hgnc_curated_tfs.txt,” motifs “motifs-v10nr_clust-nr.hgnc-m0.001-o0.0.tbl,” and the motif ranking databases “hg38_500bp_up_100bp_down_full_tx_v10_clust.genes_vs_motifs.rankings.feather” used for the pipeline was obtained from https://resources.aertslab.org/cistarget/, http://github.com/aertslab/pySCENIC/blob/master/resources/hs_hgnc_curated_tfs.txt, and https://resources.aertslab.org/cistarget/motif2tf/. As indicated in the file name, our motif search range is 500 bp upstream and 100 bp downstream of the target gene TSS. The output regulons from the cisTarget are integrated into the NicheNet’s network. Briefly, these regulons are considered statistically significant. SCENIC calculates the enrichment of a TF’s motif in its potential target genes from their gene v.s. motifs ranking matrix. First, by default, cisTarget keep the top 75% of the regulon members (target genes) of a motif by the motif’s enrichment in the gene’s promoter region. Second, a recovery curve for the motif is established by the percentage of inferred target genes recovered from parsing down the ranked list. This area under the curve (AUC) will be normalized over the AUC of all motifs to recover the same set of target genes (i.e., the inferred target genes of the motif) from all genes to obtain the Normalized Enrichment Score (NES), and a motif with NES > 3 by default is considered as significant to regulate the inferred target set. For the ChIP-seq verification of the SCENIC prediction, the chromosome coordinates were obtained from Ensembl “Homo_sapiens.GRCh38.113.chr.gtf.gz.” The ChIP-seq data was obtained from ChIP-atlas’ “Homo Sapiens ChIP: TFs and others” category, with “T cells.” “Th1 Cells,” “Th17 Cells,” “CD8+ T cells,” “memory T cells,” “Natural Killer T cells,” “Th2 Cells,” “CD4+ T cells,” and “Natural Killer cells” being selected as cell types. GenomicRanges was used to simply find the overlap between the +/- 5kb area around the target gene TSS and the peak of the TFs which are in both SCENIC predictions and the ChIP database. The percentage of a TF’s target genes being able to find at least 1 peak of that TF around its TSS in all obtained ChIP-seq data was visualized. Regulon specificity score of TF in cell subsets is calculated using the pySCENIC function “plot_rss.”

### NicheNet

NicheNet^23^ (v2.2.0) is used for ligand inference. The integrated ligand-receptor pairs, signaling network, and gene regulatory networks (prefix 21122021) were obtained from the package’s repository. TF-target gene pairs that are not in NicheNet’s original network and γδ T cell-specific ligand-receptor pairs (CD247 and its ligands EPHA2, EPCR, BTN2A1, BTNL3, BTNL8, CD1B, CD1D, MR1, CD1A, CD1C) were added to the networks with unoptimized weights of 1 given the relatively small amount of edges being added and the original author’s comments on the robustness of NicheNet without weights optimization. Functions “construct_weighted_networks,” “apply_hub_corrections,” “construct_ligand_target_matrix” from the NicheNet package were used sequentially to construct the new ligand-target matrix following NicheNet’s user guide with default parameters. The hyperparameters used during construction was the same as that of the unmodified NicheNet, which was obtained from the package’s repository under the name “hyperparameter_list.rda.” Function “evaluate_model” from the NicheNet package was used to evaluate both unmodified ligand-target matrix and new ligand-target matrix’s performance on the “gold standard” dataset that NicheNet authors used.

The application of the NicheNet is similar to and adapted from our previous work^84^. NicheNet was supplied with DEGs and Geneformer. DEGs were calculated by using Scanpy’s tl.rank_genes_groups, Wilcoxon method. DEGs with adjusted p-values < 0.05 and log_2_ fold changes greater than 1 were used as input for NicheNet. Genes expressed in at least 1% of the cells in the receiver subsets (goal states, Teff in the TRM to Teff transition and Tex in the Teff to Tex transition) were considered as background. Potential ligands were defined as ligands of NicheNet-documented receptors whose encoding genes were expressed in at least 1% of the cells. The top 15 ligands with an AUROC > 0.5 were reported in the main figure, while the top 30 were used for quantifying their enrichment in neighboring cell types. If fewer than 15 ligands met the AUROC > 0.5 threshold, all qualifying ligands were visualized and included in the enrichment analysis.

CRC TME cell type profiles were obtained from Joanito et al.^36^. The datasets, epithelial and nonepithelial, had already been quality controlled by the providers. Basic pre-processing including normalization and log transformation were done by using Scanpy’s normalize_total and log1p function. Cells labeled “LymphNode”, “Normal” were excluded. AUC calculations were performed after averaging gene expression by cell type, which is annotated by the original authors. A gene was included in the averaged profile only if its average expression was not less than 0.001 to reduce noises. For each cell type, we iterated down the ranked gene list (ranked by their average expression in the given cell type, descending order) to recover NicheNet-inferred ligands, stopping when encountering a zero-expression gene. The final area under the curve was computed using the auc function from sklearn.metrics.

### Geneformer

Geneformer^24^ (v0.1.0) was used to perform in silico overexpression. Data that only contained TRM and Teff was tokenized using Geneformer’s TranscriptomeTokenizer, retaining protein-coding and miRNA genes by default. The pre-trained model “gf-12L-95M-i4096” was fine-tuned by an input data cell type label classification task, using function “Classifier.validate” with 100 hyperparameter optimization trials, with a train-valid-test split of 0.6-0.2-0.2, max_ncells=None, freeze_layers = 6. The model with the highest accuracy was selected, and its performance as a classifier was visualized using function “plot_conf_mat.” The initial state (TRM) and the goal state (Teff) were represented by the exact mean **[CLS]** token embeddings of cells in each group. In silico overexpression was performed using function “InSilicoPerturber” with max_ncells=5000 and emb_layer=-1. The statistical test of the perturbation result is computed using function “InSilicoPerturberStats” under the mode “goal_state_shift.” Briefly, in Geneformer, significance was assessed using Wilcoxon rank-sum tests, comparing shifts induced by each gene against the background distribution of all perturbations, with Benjamini-Hochberg correction. Genes with adjusted *p* < 0.05 is considered statistically significant, and a mean cosine shift > 0.003 (empirically selected to reduce background noise) were prioritized. Genes with more than 50 detections in the initial state cells (N_Detections > 50) were reported.

## Acknowledgments

1. R. R. and D. K. B are supported by an award from Good Ventures Open Philanthropy, as well as start-up funds from Case Western Reserve University and University Hospitals.

## Funding

Open Philanthropy (DKB)

Case Western Reserve University and University Hospitals

## Author contributions

Conceptualization: RR, DKB

Data curation: RR

Formal analysis: RR Funding acquisition: DKB

Investigation: RR

Methodology: RR, DKB

Project administration: DKB

Resources: RR

Software: RR

Supervision: DKB

Validation: RR, DKB

Writing - original draft: RR

Writing - review & editing: RR, DKB

## Data and materials availability

Scripts used for this analysis can be accessed at https://github.com/Brubaker-Lab/CRCgdTCompute. All data necessary to reproduce the findings of this manuscript are available within their originally published studies as cited in the text.

