## Supplemental Figure for "A Ligand-Centered Framework for γδ T Cell Activation in Colorectal Cancer Revealed by Single-Cell and Transformer-Based Perturbation"

**Fig. S1**

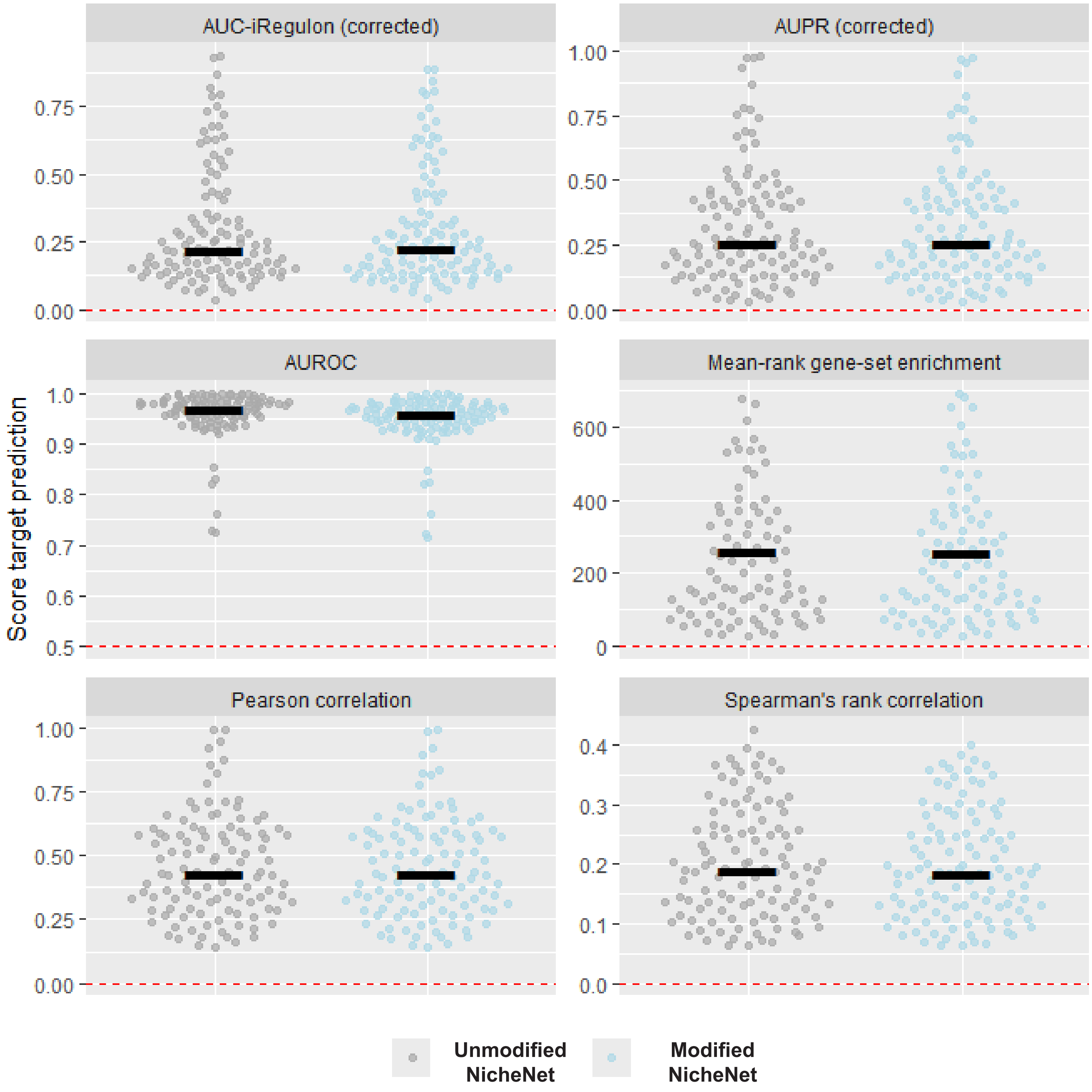


**Figure S1. Swarm plots showing the performance of the unmodified NicheNet and customized NicheNet, using the 6 metrics selected by NicheNet authors.**

Fig. S2

**a** **NicheNet-inferred top ligand's regulaotry power on top target genes**

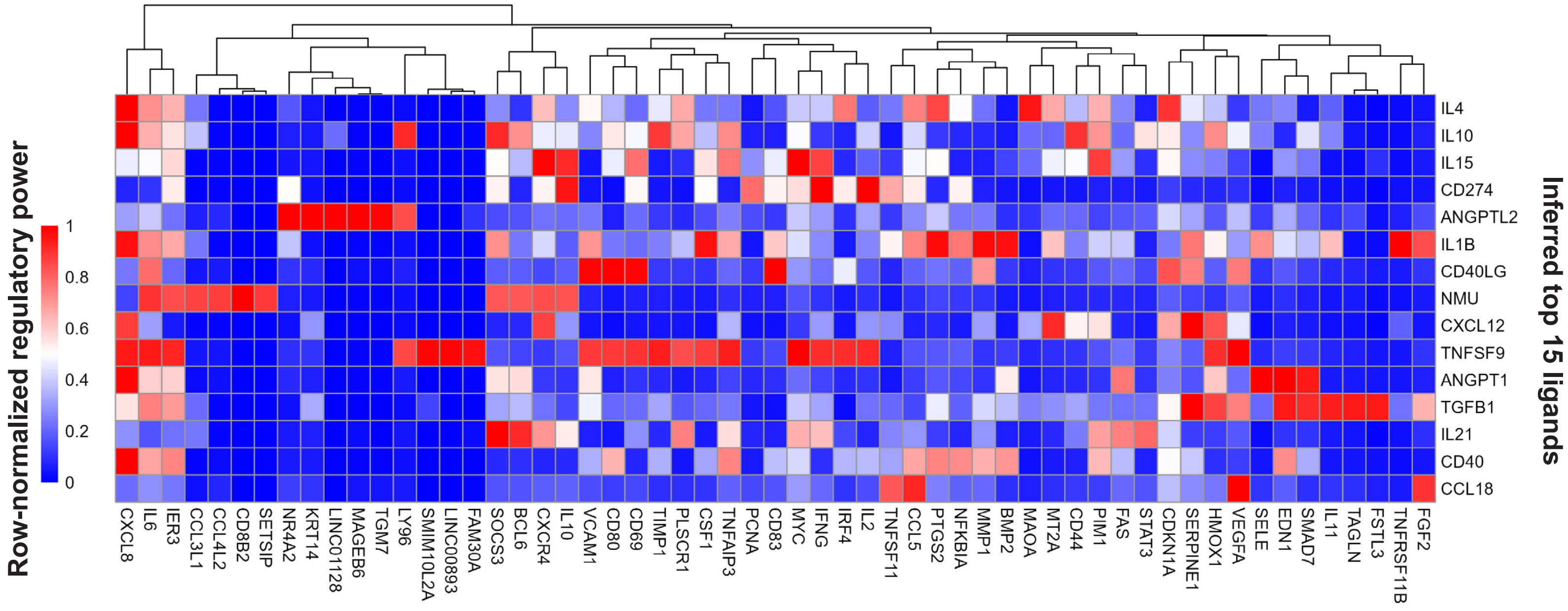


**b** **NicheNet-inferred top ligand's regulaotry power on top target genes**

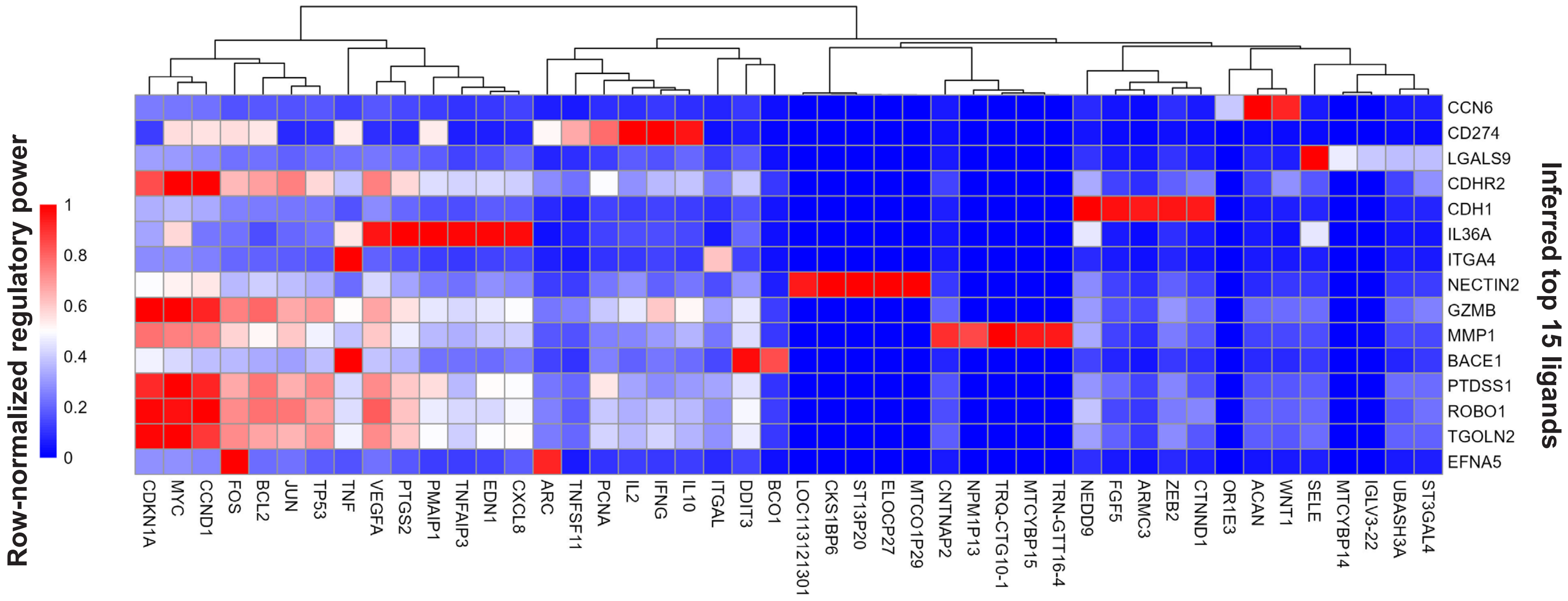


**Figure S2. Heatmaps visualizing the customized NicheNet's ligand to target gene regulatory power matrix. Rows: NicheNet-predicted top 15 ligands. Columns: Collection of top ligands' top 5 target genes. (A) TRM to Teff (B) Teff to Tex.**

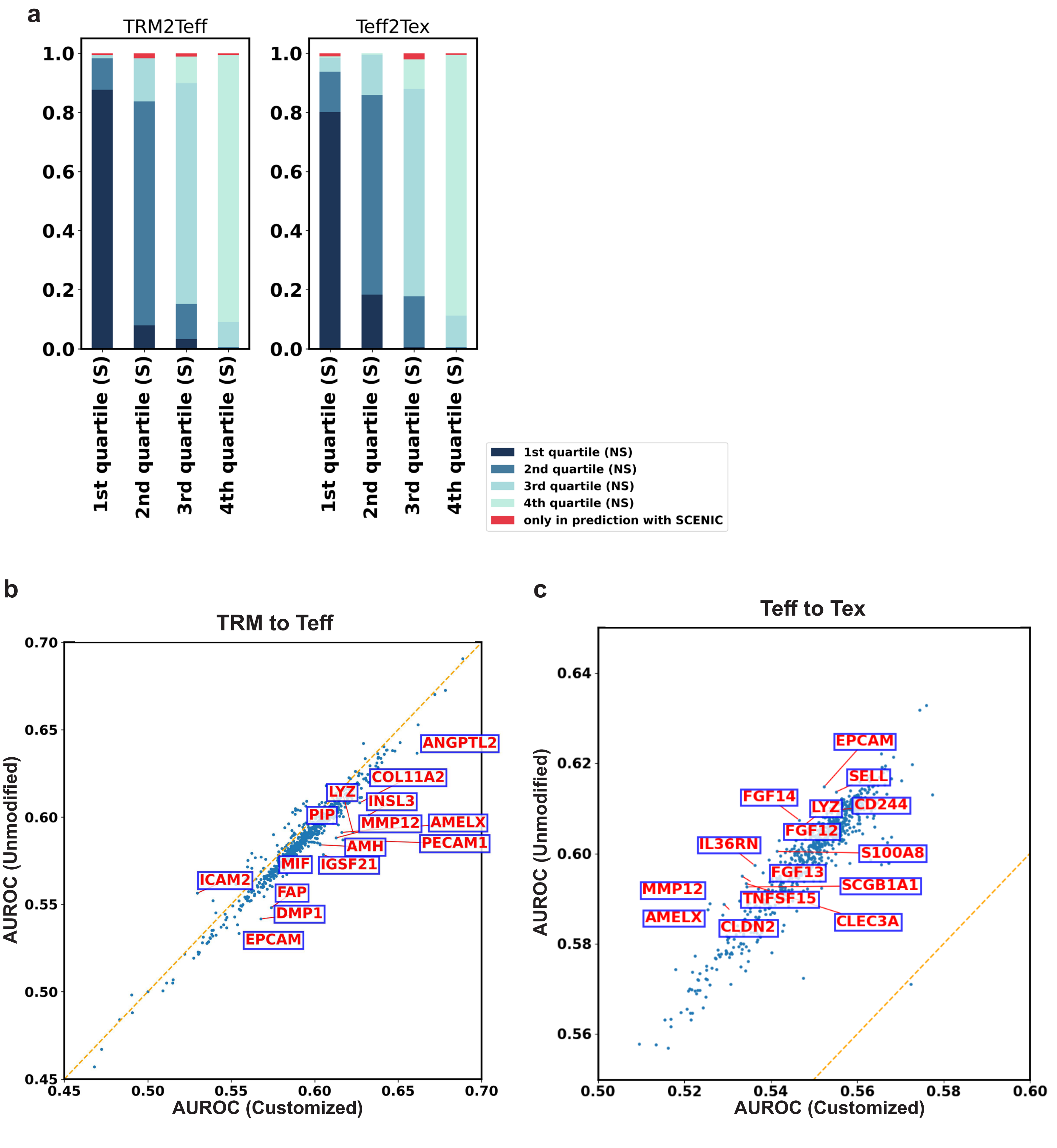


**Figure S3. Comparison of ligand prediction results between unmodified NicheNet and customized NicheNet. (A) Stacked bar plots showing the distribution of a ligand's quartile in unmodified NicheNet's results (denoted by NS) across the 4 quartiles of customized NicheNet's results. (B) Scatter plot showing the AUROC of ligands that are presents in both unmodified and customized NicheNet's results for TRM to Teff. (C) Scatter plot showing the AUROC of ligands that are presents in both unmodified and customized NicheNet's results for Teff to Tex.**

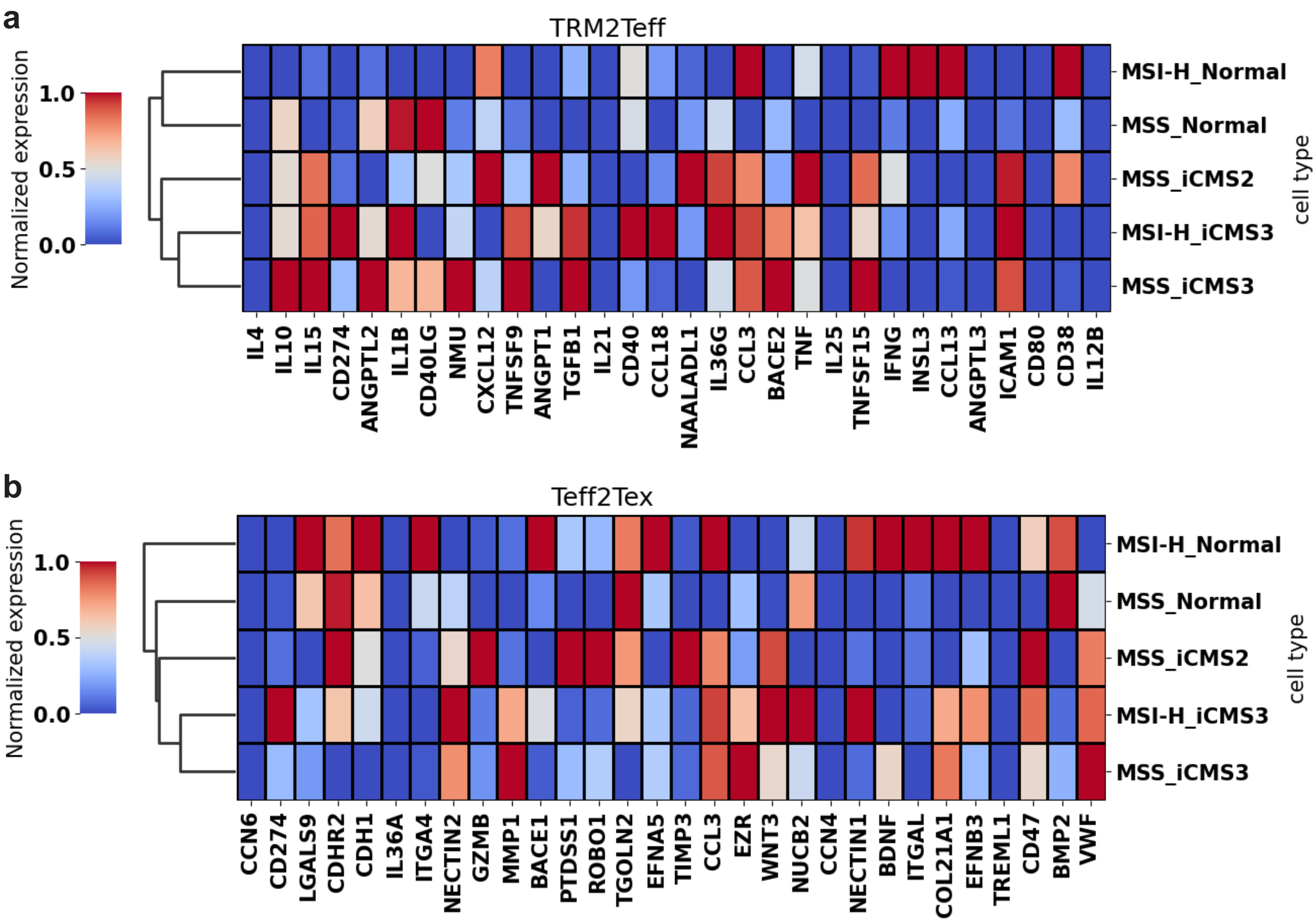


**Figure S4. Heatmap showing the expression of top 30 NicheNet-inferred ligands in CRC epithelial cell types. (A) TRM to Teff ligands enrichment. (B) Teff to Tex ligands enrichment.**

**Fig. S5**

**a**

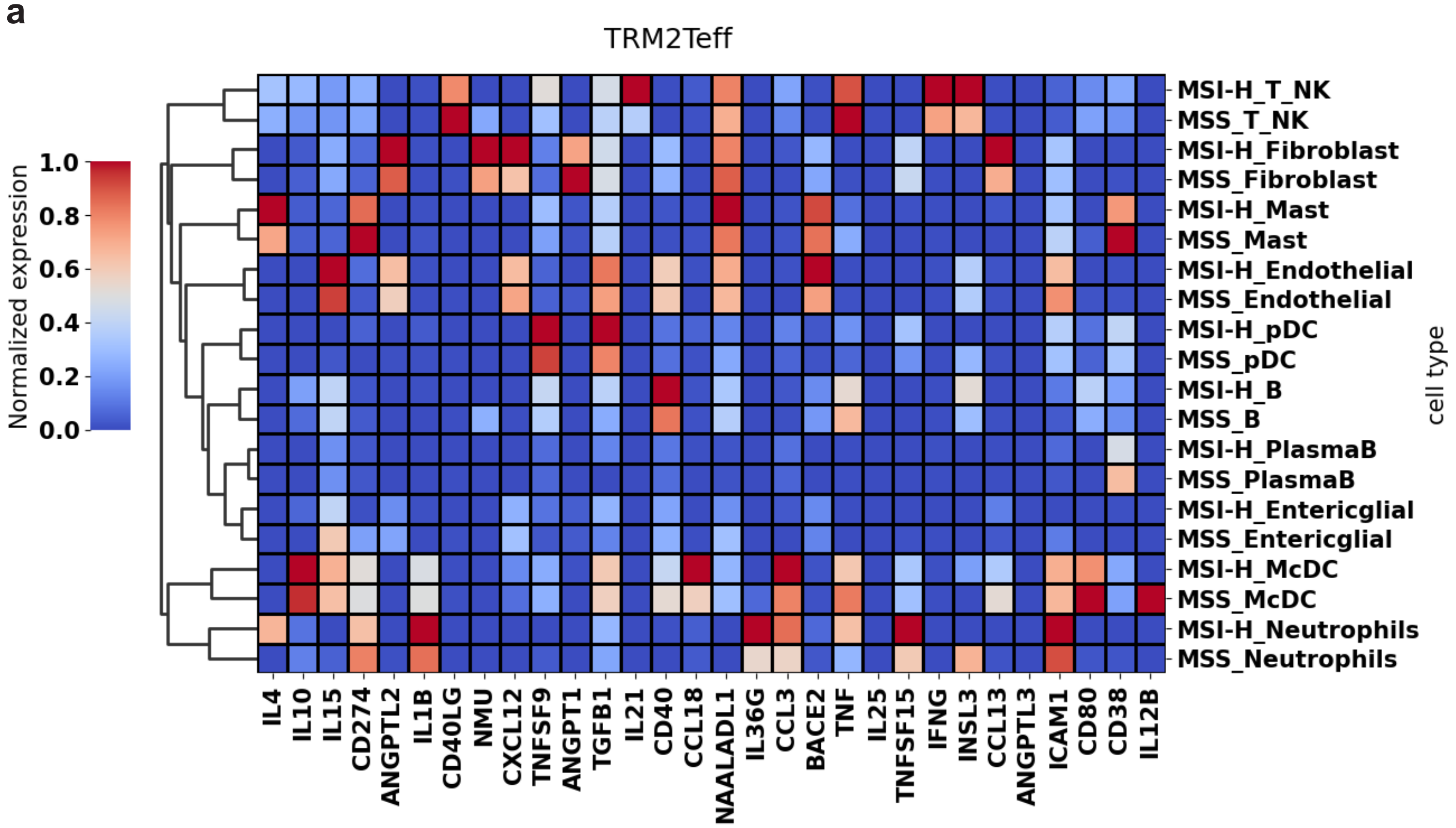


**b**

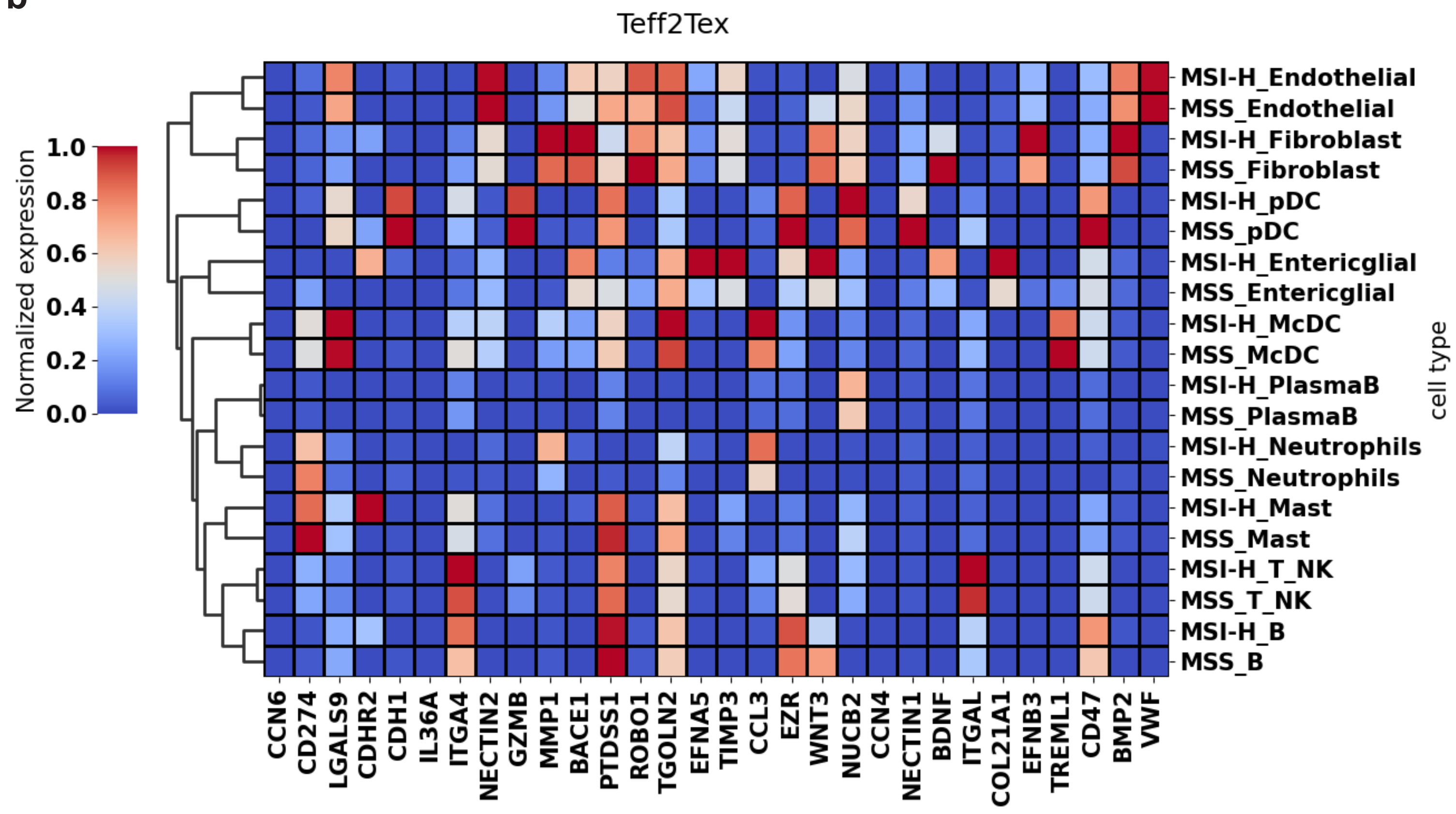


**Figure S5. Heatmap showing the expression of top 30 NicheNet-inferred ligands in CRC non-epithelial cell types. (A) TRM to Teff ligands enrichment. (B) Teff to Tex ligands enrichment.**

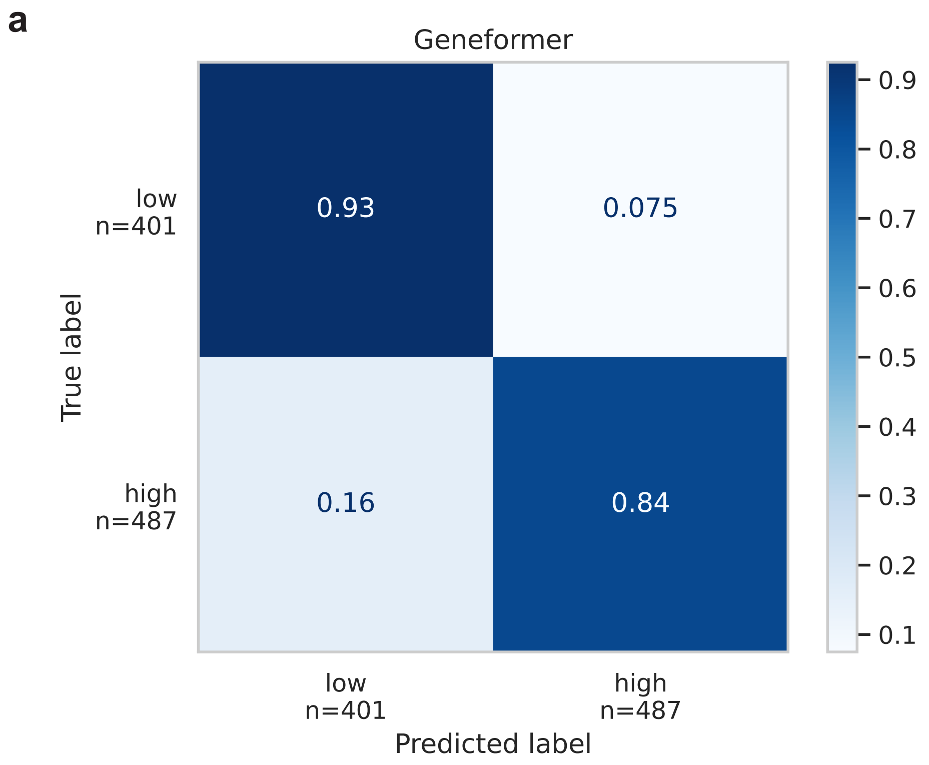


**Figure S6. Heatmap showing the fine-tune Geneformer's performance in classifying TRM and Teff cells in the input data.**
